# *T* and *Z*, Partial Seed Coat Patterning Genes in Common Bean, Provide Insight into the Structure and Protein Interactions of a Plant MBW Complex

**DOI:** 10.1101/2023.09.28.560026

**Authors:** Phillip E. McClean, Jayanta Roy, Christopher L. Colbert, Caroline Osborne, Rian Lee, Phillip N. Miklas, Juan M. Osorno

**Affiliations:** Department of Plant Sciences, North Dakota State University, Fargo, North Dakota 58108 USA; Genomics and Bioinformatics Program, North Dakota State University, Fargo, North Dakota 58108 USA; Department of Chemistry and Biochemistry, North Dakota State University, Fargo, North Dakota 58108 USA; USDA-ARS, Grain Legume Genetics Physiology Research Unit, 24106 N. Bunn Rd., Prosser, Washington, USA

**Keywords:** AlphaFold2, AlphaFold-Multimer, bHLH, common bean, MBW, MYB, protein interactions, protein modeling, WDR

## Abstract

Flavonoids are secondary metabolites associated with seed and flower color. A ternary MBW protein complex consisting of interfacing **<u>M</u>**YB, **<u>b</u>**eta-helix-loop-helix (bHLH), and **<u>W</u>**D40 repeat (WDR) proteins controls the expression of late biosynthetic genes in the flavonoid pathway. *P*, the master regulator gene of flavonoid expression in common bean (*Phaseolus vulgaris* L.) was recently determined to encode a bHLH protein. Two other genes, *T* and *Z*, are also historically considered regulators since they determine the distribution of color in seeds and flowers. These two genes were cloned using reverse genetics based on genetic mapping, candidate gene selection, phylogenetic analysis, and mutant analysis. Domain and AlphaFold2 structure analyses determined *T* encoded a seven-bladed β-propeller WDR protein, while *Z* encoded a R2R3 MYB protein. Modeling of the Z MYB/P bHLH/T WDR MBW complex identified interfacing sequence domains and motif in all three genes that are conserved in dicots. One Z MYB motif is a possible β-MoRF that only appears in a structured state when Z MYB is modeled in a MBW complex. Complexes containing mutant T and Z proteins changed the interaction of members of the complex in ways that would alter their role in regulating expression of genes in the flavonoid pathway.

**Highlight:** The MBW complex of bean comprised of the classic *Z*, *P*, and *T* genes associated with seed coat patterning defines critical proteins interactions for flavonoid gene expression.

## Introduction

The seed coat of many plant species is commonly solid colored. The color is generally the result of the flavonoid biosynthetic pathway. Common bean is somewhat unique since its seed coat can be either solid colored, patterned with a light background splashed with stripes or mottles, or partially colored with white and colored zones. The partial seed coat types intrigued early breeders and quickly became a research focus following the rediscovery of Mendel’s research. The master regulator of all patterns and colors is the *P* (*=Pigment*; Emerson 1909) gene. Any genotype with the homozygous recessive *pp* genotype does not express any color in the seed coat (or flower). All partial seed coat patterns require, in addition to the dominant *P* allele, a homozygous recessive *tt* genotype at the *T* (*=Total*; Emerson 1909) gene. The extent of the darker region is controlled by the interaction of the *tt* genotype and various alleles at the *Z* (=*Zonal*; Tschermak 1912), *Bip* (=*Bipunctata*; Lamprecht 1932), *J* (=*Joker*; Lamprecht 1932), and *Fib* (=*fibula*; Bassett 2001) genes. For example, *P t Z Bip J fib* genotypes express full seed color, *P t z Bip J fib* express dorsal seed color (virgarcus pattern), and *P t z bip J* express color in two regions neighboring the hilum (bipuncta pattern). *Z* and *J* also control the expression of color in the hilum ring, a layer of cells bordering the hilum. In a *P T z j* background, color is not expressed in the hilum ring (see a detailed summary in Bassett 2007). This led to the suggestion that many if n ot all of these genes are regulators on the flavonoid pathway.

The *P* gene, first gene to be described as a molecular component of color gene regulation in bean, encodes an ortholog of other **b**eta-helix-loop-helix proteins (bHLH; McClean *et al*. 2018) that functions in a ternary complex along with a **W**D40 repeat (WDR) protein and one of several **M**YB proteins. This complex, termed MBW, regulates the late biosynthetic genes (LBGs) of the flavonoid pathway (Stracke *et al*. 2007; Stracke *et al*. 2010). The WDR protein is encoded by a single gene in most species, one or a few genes encode the bHLH protein, and one of multiple MYB activators or repressors interact with its two complex partners to fine tune expression of the flavonoid LBGs. The MBW complex involves several protein interactions. The MYB and bHLH proteins were initially shown to physically interact (Goff *et al*. 1992) and bind upstream of the flavonoid LBGs pathway (Goff *et al*. 1990). The bHLH and WDR proteins interact and form a ternary complex with a MYB factor (Payne *et al*. 2000; Zhang *et al*. 2003). WDR and MYB factors were reported to interact in some (Cui *et al*. 2021; Liu *et al*. 2018) but not all (Gao *et al*. 2018) species. Mutations of any member of the MBW complex directly affect the expression of the flavonoid pathway LBGs and alters seed coat flavonoid composition. For example, Arabidopsis mutants of the MBW MYB gene (*TT2*; Nesi *et al*. 2001) have golden yellow testa, while mutants of the bHLH (*TT8*; Nesi *et al*. 2000), and WDR (*TTG1*; Walker *et al*. 1999) have colorless testa. In a similar vein, mutants of the common bean *P* gene, the common bean ortholog of the Arabidopsis bHLH *TT8* gene, have colorless seed coats (McClean *et al*. 2018). MBW genes from other species also have a strong effect on seed coat color expression. Three legume genes, pea *A2* (Hellens *et al*. 2010), *Medicago truncatula* MtWD40-1 (Pang *et al*. 2009; Meng *et al*. 2019), and *Vicia faba zt1* (Gutierrez and Torres 2019), encode WDR proteins, and mutants of these genes express white flowers and seeds.

Because of their apparent regulatory function, the focus here is to identify candidate genes for *T* and *Z* in common bean. *T* was selected since a mutant allele of this gene is the master regulator of all partial seed coat phenotype suggesting a role in flavonoid biosynthesis. *Z* was a focus because *tt* mutants further restrict color expression beyond that of *tt* genotypes. Since these mutants result in zones without color expression, and that a functioning MBW complex is required for color expression, the further assumption was that *T* and *Z* function as regulators of flavonoids and possibly components of the MBW complex. Here, genetic analysis, candidate gene identification, phylogenetic comparisons, and protein modeling identified a β-propeller, WDR-encoding gene as a strong candidate gene for the*T* gene. Using the same experimental approach, a MYB-encoding gene model was identified as a *Z* candidate. To develop a view of how these two proteins might physically interact as part of bean MBW complex, wild type and mutant versions of the *T*, *Z*, and *P* genes were modeled by the recently released AlphaFold2 (AF2) protein modeling algorithms (Jumper et al. 2021) to consider the structural effects of the mutants. Finally, modeling dimers and trimers of the T, Z, and P proteins for the first-time defined specific interacting domains among the three wild type proteins and the effects mutant proteins on those interactions.

## Material and Methods

### Plant material

5-593 (PI 608674) is a genotype selected by Dr. Mark Bassett (University of Florida) from a genetic stock developed by Dr. Jim Beaver (University of Puerto Rico). 5-593 is dominant for nearly all seed coat color and pattern genes. The 5-593 genotype for the patterning and color genes is: *T Z Bip P* [*C r*] *J G B V Rk Gy sal.* Dr. Bassett introgressed recessive alleles for many color and pattern genes into 5-593 through a backcrossing program to the level of BC_3_. Candidate recessive *t* and/or *z* alleles were sequenced from the following donor lines: 65-73 (PI 451802), homozygous recessive for a t allele; V0869 (PI 527806) and V0919 (PI 527820), homozygous for recessive z; and Earliwax (PI 549618), homozygous recessive for both *t* and *z* (Fig. 1; Table 1). Dr. Bassett’s genetic marker introgression lines and donor genotypes for the mutant alleles in the introgression lines were evaluated for sequence variability with *T* and *Z* PCR Allele Competitive Extension (PACE) markers for the two genes (Supplementary Table S1). The two PACE markers were also used to determine the allelic state of these two genes in a wide collection of genotypes from the Middle America Diversity Panel [MDP (Moghaddam *et al*. 2016; Supplementary Table S2] and Andean Diversity Panel [ADP (Cichy *et al*. 2015); Supplementary Table S3].

**Fig. 1.**
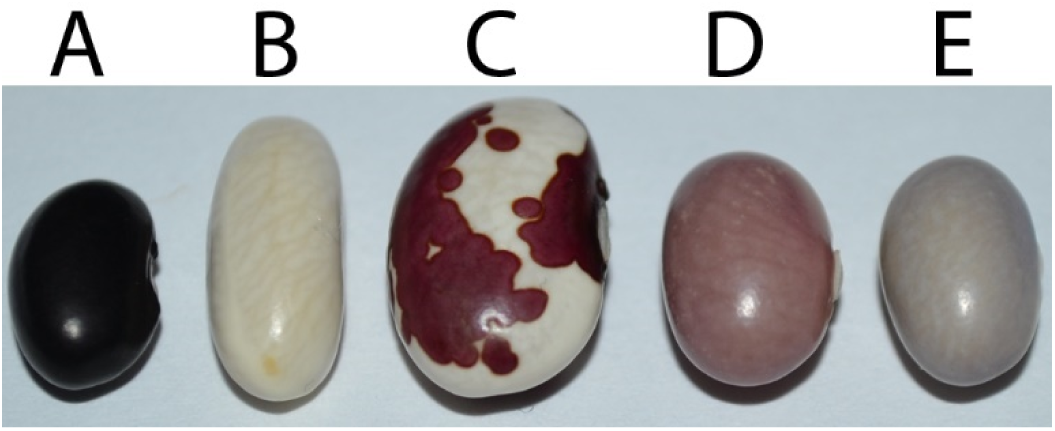
Germplasm used to map the *T* and *Z* genes of common bean. The *T* and *Z* alleles for each genotype are in parenthesis. (A) 5-593 (*T Z*). (B) Earliwax (*t*^EW^ *z*^EW^). (C) 65-73 (*t*^65-73^ *Z*). (D) V0869 (*T z*^EW^). (E) V0919 (*T z*^EW^).

### DNA isolation, amplicon sequencing, and CDS assembly

DNA was extracted from young leaf tissue using the Mag-Bind Plant DNA Plus kit (Omega Bio-Tek; https://www.omegabiotek.com/product/mag-bind-plant-dna-plus-96-kit/). DNA amplicons were generated by PCR amplification with the primers listed in Supplemental Table S4. The reaction was performed in a 25-µl volume over 45 cycles using the annealing temperatures specific for each pair of primers. The NEB Monarch Gel Extraction Kit (https://www.neb.com/products/t1020-monarch-dna-gel-extraction-kit) was used to isolate individual amplicon fragments, and individual fragments were sequenced by Eton Bioscience Inc. (https://www.etonbio.com/). The coding region of the fragment sequences were assembled using the Staden Package (https://sourceforge.net/projects/staden/; Staden 1996) into the CDS sequence.

### Phylogenetic tree building and T and Z gene alignment of orthologs

A neighbor-joining (NJ) phylogenetic tree consisting of proteins homologous to the *T* and representing the breadth of Angiosperm diversity are listed in. These proteins (Supplementary Table S7) were downloaded from Phytozome 13 (https://phytozome-next.jgi.doe.gov/). The proteins were identified via blastp search in Phytozome using the Pv5-593.09G047300 protein sequence as the query. Functionally verified flavonoid activator and repressor MYB proteins identified from the literature (Supplementary Table S8) were used to construct a NJ tree that included the *Z* gene model. Each NJ tree was built using MEGA 7 (Kumar *et al*., 2016). Tree building was based on the MUSCLE protein alignment algorithm (Edgar, 2004). The Jones-Taylor-Thornton substitution model was used for tree development. The NJ tree was obtained using 1000 bootstrap replicates. Protein alignment was performed using the T-Coffee server (http://tcoffee.crg.cat/apps/tcoffee/do:regular), and the alignments were colored using MS Word.

### PCR Allele Competitive Extension marker development

PACE markers were developed for the candidate *T* gene mutations discovered in Earliwax (1-bp deletion) and 65-73 (39-bp deletion). The nucleotide deletion in Earliwax is at position Pv09:10,713,827. The 39-bp deletion 65-73 begins at position PV09:10,713,103 bp. PACE markers were also developed for two Earliwax mutations in the*Z* gene candidate. One mutation was a SNP at position Pv03:34,523,093 bp, and the second mutant was a deletion in the interval: Pv03:34,522,869..Pv03:34,522,880 bp. Each position is relative to the 5-593 reference genome assembly (https://phytozome-next.jgi.doe.gov/info/Pvulgaris5_593_v1_1). For each polymorphism, the unique forward and common reverse primers (Supplemental Table 5) were designed by the 3CR Bioscience (https://3crbio.com/) primer development service. The genotyping reactions contained 2 µl (of approximately 10 ng/ul) of DNA, 0.15 µl of primer mix (12 µM of each allele-specific forward primer, 30 µM of the common reverse primer), 4 µl of 3CR Bioscience PACE Master Mix Standard Rox, and 1.85 µl H_2_O for a total volume of 8 µl. PCR amplifications were performed with the following PCR cycling conditions: 94 °C for 15 min, followed by a touchdown profile of 10 cycles at 94°C for 20 s, and 65°C for 1 min with a 0.8°C reduction per cycle, followed by 40 cycles at 94°C for 20 s and 57°C for 1 min. End point reads for the 96 well plates were collected using the Bio-Rad CFX96 Touch Real-Time PCR Detection System to determine the allele of each genotype. Readings were made after PCR, and the relative fluorescent unit (RFU) data was output using the Bio-Rad CFX Maestro 2.0 software.

### Genetic mapping of the T WDR and Z MYB candidate genes and variation in common bean germplasm

F_2_ populations (Brady *et al*. 1998; Bassett *et al*. 1999), based on crosses of genetic stocks developed by Dr. Bassett (Univ. of Florida) were used to map candidate genes relative to the*T* and *Z* phenotypes. PACE markers were developed that span the interval defined by the physical position of OM19_400_ and AM10_560_, two RAPD markers determined by Brady *et al*. (1998) to be linked to *T* and *Z*, respectively. One F_2_ population (4-68) was used to map *T* with the new markers and its relationship to the original *T* marker. The *Z* gene was mapped relative to the *Z* marker position in three F_2_ populations (4-76, 6-273, and 6-160.161). The cross, donor recessive allele, and phenotypic ratios can be found in Supplementary Table S6.

### Protein sequence analysis and structure predictions of the bHLH P protein, the WDR T gene protein and the MYB Z gene protein

The domain structure of the Pv5-593.09G047300 and Pv5-593.03G127600 proteins was determined with the SMART application (http://smart.embl-heidelberg.de/) as implemented in Interpro (https://www.ebi.ac.uk/interpro/). Intrinsically disordered regions (IDR) of the P bHLH, Z MYB, T WDR proteins were determined using the IUPred3 (Erdős *et al*. 2021) and DISOPRED3 (Jones and Cozzeto 2015) algorithms. Regions with disorder scores greater than 0.5 were considered to be disordered.

For protein structure predictions, the AlphFold2 (AF2; Jumper *et al*. 2021) neural network, as implemented in ColabFold (Mirdita *et al*. 2022; https://github.com/sokrypton/ColabFold), was used for all protein structural predictions. Protein complex predictions were generated using Alphafold-Multimer (AF-Multimer; Evans *et al*. 2022), also as implemented in ColabFold. MMseqs2 (Steinegger and Söding 2017) was used for the homolog search against the UniRef90 database. Five models were developed, and the model with the highest predictive-local distance difference test (pLDDT) score was selected as the best model. The predicted protein structure was visualized using the ChimeraX (Pettersen*et al*. 2021). The structural interaction of the modeled bean MBW complex was analyzed with the ‘Protein interfaces, surfaces and assemblies’ (PISA) service at the European Bioinformatics Institute (https://www.ebi.ac.uk/pdbe/pisa/; Krissinel and Henrick 2007). The ChimeraX Matchmaker function was used to calculate the least-squares-fit root-mean-square deviation (RMSD) between any two predicted structure models.

## Results and Discussion

### Identifying T and Z candidate genes

*T* cosegregates with marker OM19_400_, while *Z* is closely linked with marker AM10_560_ (Brady *et al*. 1998). These genes were subsequently mapped to chromosomes Pv09 and Pv03, respectively (McClean *et al*. 2002). Cloning, physical and genetic mapping placed the *T* gene in the 10.11-11.53 Mb interval on chromosome Pv09 (Fig. 2A), while *Z* mapped to a 33.70-35.11 Mb interval on Pv03 (Fig. 2B).

**Fig. 2.**
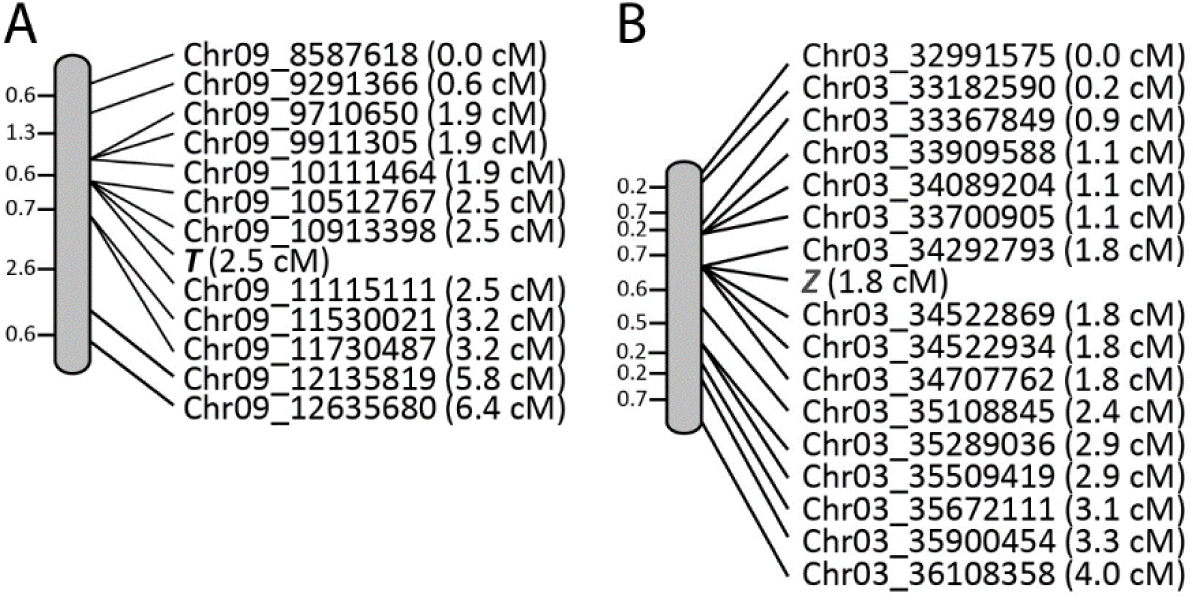
Genetic maps of *T* and *Z* developed using PACE SNP markers bordering the position of RAPD markers linked to each gene. (A) The *T* gene was mapped in population 4-68 (Brady *et al*. 1998). (B) The *Z* gene was mapped in three populations (4-76, 6-273, and 6-160.161). For both (A) and (B), the physical position of the polymorphic SNP follows the chromosome designation, and the genetic position is located in parenthesis.

Based on the strong phenotypic effects of *T* and *Z* on color expression in seed coats and flowers, these physical intervals were searched for gene models in the 5-593 reference genome that enzymatic or regulatory genes of the flavonoid pathway as described by Wu *et al*. (2018). Only one gene (*Pv5-593.09G046900*), which met this criteria encodes a flavonol 3-O-methyltransferase (E2.1.1.76) that decorates the three position of flavonols with a methyl group, was located within the *T* gene interval. Its protein sequence from Earliwax (PI 549618) and 65-73 (PI 451801), that carry a recessive *t* allele, was identical to the dominant *T* allele of 5-593, this gene model was not considered further as a *T* gene candidate.

Based on the catalog of Arabidopsis MYB, βHLH, and WDR transcription factors described by Zhang and Schrader (2017), a *T* gene candidate, *Pv5-593.09G047300*, that encodes a WDR protein begins at position 10,712,956 bp of the 5-593 reference genome sequence was located within the *T* genetic interval (Fig. 2a). Based on replicated RNA-seq data for 5-593 available at Phytozome, the gene was expressed in four progressive stages of seed coat development. A blastp analysis found *Pv5-593.09G047300* was orthologous to the Arabidopsis *TTG1* gene (AT5G24520) that also encodes a WDR protein (Walker *et al*. 1998) that anchors the TT8 βHLH in the MBW complex. Pv5-593.09G047300 is also orthologous to the *Medicago truncatula* MtWD40-1 WDR protein (Medtr3g092840) that controls seed coat color expression (Pang *et al*. 2009), the *Pisum sativum* A2 WDR protein (NCBI identifier ADQ27318.1) that controls flower color (Hellens *et al*. 2010), and *Glycine max* GmWD40 (Glyma.06G136900) that is required to activate proanthocyanin production (Lu *et al*. 2021). MtWD40-1 was recently confirmed as the WD40 component of the *M. truncatuala* MBW complex (Meng *et al*. 2019). A multi-sequence alignment found the sequence of these five proteins to be highly conserved (Supplementary Fig. S1).

Gene model *Pv5-593.03G127600,* which encodes a R2R3-MYB protein, was identified as a candidate for the *Z* gene using the same approach as with *T*. Replicated RNA-seq analysis found the gene to be expressed only during the four stages of seed coat development. *Pv5-593.03G127600* was orthologous to the Arabidopsis TT2 protein (Nesi *et al*. 2001), one of the several MYB factors that functions in the Arabidopsis MBW complex. *Pv5-593.03G127600* was also orthologous to the *M. truncatula* MtMYB14 protein that interacts with another R2R3-MYB protein, MtMYB5, to activate LBGs of the flavonoid pathway (Liu *et al*. 2014). This activation is in conjunction with MBW proteins MtTT8 (a bHLH ortholog of the Arabidopsis TT8 protein) and MtWD40-1 (a WDR ortholog of Arabidopsis TTG1 protein). The blastp analysis also identified two orthologs in the Williams82 reference soybean genome, Glyma.13G109100 and Glyma.17G050500 that were labeled GmTT2A and GmTT2B, respectively (Lu *et al*. 2021). These proteins interact with the soybean bHLH and WDR proteins to activate expression of flavonoid LBGs (Lu *et al*. 2021). Overexpression of *Glyma.17G050500* in transgenic soybeans resulted in greater flavonoid pigmentation in the hilum. Similarly, *Z* also controls hilum pigmentation in common bean (Bassett *et al*. 1999). The sequence alignment of the candidate Z protein, along with the AtTT2, MtMYB14, and GmTT2A proteins, found a high degree of conservation within the N-terminal R2 and R3 MYB domains while the C terminal region was much less conserved (Supplementary Fig. S2).

### Phylogenetic analyses of T WDR and Z MYB and their orthologs

A blastp analysis across the genus *Phaseolus* and 22 other species representing the taxonomic breadth of the Angiosperms was performed. The T WDR (Pv5-593.09G047300) protein sequence was identical to its orthologs from the other three bean reference genome annotations (G19833 v2.1, UI111 v1.1, and Labor Ovalle v.1.1) and tepary bean (*P. acutifolius*). T WDR was 98% and 93% identical to its cowpea (*Vigna unguiculata*) and soybean (*G. max*) orthologs, respectively. The percent amino acid identity for the remainder of the species ranged from 84% (*Cicer arietinum* and *M. truncatulata*) to 62% (*Panicum hallii* and *Sorghum bicolor*). A phylogenetic tree (Fig. 2A) found these identity values reflected the taxonomic distance from common bean (Supplementary Table S7). At the taxonomic order level, the NJ treeagrees with the current angiosperm tree (APG IV, 2016). Among the Fabales species, the gene tree is consistent with the most recent legume species tree based on multi-locus sequence data (Koenen *et al.,* 2020).

Plant MYB proteins typically contain two conserved N terminal MYB domains, R2 and R3, and the R3 domain interacts with bHLH proteins. The variable C-terminal portion of the proteins is thought to provide the distinct functionality of each member of the family (Millard *et al*. 2019). These functions include anthocyanin and proanthocyanin biosynthesis, abiotic and biotic stress responses, cell fate, meristem and lateral organ development, hormone-induced growth, and embryogenesis (Dubos *et al*. 2010). To provide insight into the functionality of the *Z* MYB gene in color expression, the phylogeny presented here (Fig. 2B) focused on proteins experimentally shown to be associated with anthocyanin and proanthocyanin regulation. Z MYB was placed in a highly supported clade of genes previously shown to activate proanthocyanins production.

### T WDR and Z MYB polymorphisms among common bean genotypes

A blastp analysis with the T WDR sequence as a query found orthologs in the three other common bean reference genomes: *Phvul.009G044700* (G19833; Schmutz *et al*. 2014), *PvUI111.09G045300* (UI 111), and *PvLabOv.09G054700* (Labor Ovalle) were identical at the protein level. The CDS sequence of the UI 111 and Labor Ovalle Middle American orthologs were identical to the 5-593 sequence and differed from the Andean ortholog G19833 sequence by two SNPs: T102C and C471A. Based on the fact that G19833 and 5-593 carry the dominant *T* allele (Bassett *et al*. 2011), it can be concluded that UI 111 and Labor Ovalle also carry the dominant *T* allele. This is consistent with the fact that neither of these genotypes has partially colored seeds.

The *T WDR* nucleotide sequence was determined for two recessive *tt* genotypes, 65-73 and Earliwax (Fig. 1). The recessive *t* allele of the 65-73 genotype contained an in-frame 39 nucleotide deletion (CDS:25-63) that results in a protein that lacks 13 amino acids near the N-terminal region (Supplementary Fig. 3). This allele was designated *t*^65-73^. A single nucleotide deletion at CDS position 752 of the genotype Earliwax resulted in a stop codon that reduced the protein lengths by 82 amino acids relative to the Pv5-593.09G047300 protein. This allele was designated *t*^EW^. A third recessive allele was detected in PI 632734. The allele is identical to *t*^EW^ except for an additional 5-bp deletion beginning at position 426 of the CDS. The mutation introduced a premature stop codon that shortened the protein to 178 amino acids. This allele was designated *t*^PI632734^.

A blastn analysis with the *Z MYB* against the other three other common bean reference genomes found the UI 111 sequence identical to the 5-593 sequence. The G19833 and Labor Ovalle orthologs contain 12 and 18 nucleotide (nt) in-frame deletions, respectively, that shorten their protein sequences by four and six amino acids, respectively (Supplementary Fig. 4). They both share a common SNP, while Labor Ovalle contains one non-synonymous and five additional synonymous SNPs. The G19833 allele was named *Z*^G19833^, and the Labor Ovalle allele was named *Z*^LO^.

The sequence of recessive *zz* donors lines V0869 (PI527806), V0919 (PI527820), and Earliwax (PI549618) used to develop multiple introgression lines, was extracted from a 40X resequencing data set (McClean *et al*. 2022). Relative to 5-593, all three lines share the 12 nt in-frame deletion with G19833. In addition, each line contained a non-synonymous G315T SNP of the CDS sequence which led to a K105N amino acid substitution not found in G19833. This allele was designated as *z*^EW^. G19833 contains a colored hilum ring that is controlled by the dominant *Z* allele. Since it shares the 12 nt deletion with the three z donor parents, which lack the hilum ring, that deletion does not appear to be the functional mutation. Rather, the G315T SNP responsible for the K105N substitution in recessive *zz* lines appears to be the candidate causative mutation. This suggestion is further supported by the fact that the lysine at position 105 is conserved in all other functional MYB proteins associated with activators and repressors of proanthocyanin and anthocyanin synthesis described above (Fig. 2B).

### T and Z gene diversity among Middle American and Andean bean genotypes

PACE markers specific to mutations *t*^65-73^ and *t*^EW^ alleles (Supplemental Table 4) were tested on a selection of genotypes used by Dr. Bassett to create a series of introgression lines containing the recessive*t* allele (Supplemental Table 1). Each line whose *t* allele traced back to Earliwax was positive for the *t*^EW^ PACE marker, while *t* introgression lines derived from 65-73 were positive for the *t*^65-73^ PACE marker allele. Members of the MDP (n=284) and ADP (n=256) populations were also screened with these markers. All MDP (Supplemental Table 2) and ADP (Supplemental Table 3) genotypes carried the dominant*T* allele at each of the two markers. For the *Z* candidate gene, PACE markers for both the G315T SNP and the 12 nt deletion in the *z*^EW^ allele were developed. The dominant *Z* allele predominated in the MDP (93.3%), whereas a large majority of the ADP carried the recessive *z*^EW^ allele (96.8%).

### Genetic analysis of the T and Z genes

The genetic relationship between the *T* PACE markers for *Pv5-593.09G047300* and the *Z* PACE markers and *Pv5-593.03G127600* were evaluated in F_2_ populations (Supplemental Table 6). The *t*^EW^ PACE marker was found to co-segregate with flower color data from the 4-68 F_2_ population that segregates for the *t*^EW^ allele (Brady *et al*. 1998). Every purple-flowered individual (n=60) was either homozygous or heterozygous for the dominant 5-593 *T* marker allele, while all white flowered individuals (n=19) were homozygous for the recessive *t*^EW^ marker allele. This provided additional genetic evidence that the *Pv5-593.09G047300* was a strong candidate gene for the *T* gene.

The linkage between the *Z* gene and the *Pv5-593.03G127600* z^EW^ marker was tested with three populations segregating for the Earliwax *z* allele. The 4-76 population segregated 3:1 for presence (*Z*_) absence (*zz*)of the hilum ring. All F_2_ plants with a hilum ring (n=63) were either homozygous or heterozygous for the dominant 5-593 *Z* marker allele, while those without a hilum ring (n=11) contained the recessive z^EW^ marker allele. The 6-160.161 F_2_ population segregated in the expected 3:1 ratio of self-colored (*ZZ*) + ambigua (*Zz*^EW^) seed patterns versus the virgarcus (*z*^EW^*z*^EW^) pattern. All individuals in the population expressing self-colored or ambigua pattern (n=60) contained the 5-593 *Z* marker allele, while the virgarcus individuals (n=19) all contained the z^EW^ marker allele. For the third population, 6-273, the expected 3:1 ratio [self-colored (*ZZ*) + Anasazi (*Zz*^EW^) versus Anabip + virgarcus (*z*^EW^*z*^EW^)] was observed. All self-colored or Anasazi patterned plants (n=53) carried the 5-593 *Z* marker allele, and those plants expressing the Anabip or virgarcus pattern (n=19) contained the z^EW^ marker allele. The collective data across all three populations (n=225) provides compelling evidence that *Pv5-593.03G127600* is a very strong candidate gene model for the *Z* gene.

### Protein modeling of T, Z, and P

Domain analyses identified four WD-40 repeats (SMART accession number: SM00320; Pfam accession: PF00400) in the T WDR protein and its soybean (GmWD40), Medicago (MtWD40-1), and Arabidopsis (AtTTG1) orthologs (Supplementary Fig. S1). The AF2 model contained seven-blades, each consisting of four anti-parallel β-strands labeled a, b, c, and d (Smith *et al*., 1999; Jain and Pandey 2018), to form the classic toroidal β-propeller structure of WDR proteins. As an indication of AF2 model quality, the predictive score (pLDDT) for T WDR was >90 for more than 82% of the residues and >70 for more than 90% of the residues (Fig. 4A). This agrees with the DISOPRED3 and IUPre disorder analyses where only a few residues were considered disordered (Fig. 4A). The predicted structure of T WDR was in good agreement with the predicted structure of GmWD40, MtWD40-1, and AtTTG1 with C_α_ RMSD values of 0.714 Å, 1.431 Å, and 2.585 Å, respectively, across the full 336 residues.

Domain analysis of the Z MYB protein and its orthologs GmTT2B MYB, MyMYB14, and AtTT2 MYB, revealed two MYB domains (SMART accession number: SM00717; Pfam accession: PF00249). Plant MYB protein typically contain two tandem repeats with three alpha-helices each arranged in helix-turn-helix motifs that are involved in DNA binding (Feller *et al*. 2011). Structure prediction by AF2 on the Z protein candidates contained the expected six N-terminal helices, three each for the R2 and R3 MYB domains (pLDDT>90). The C-terminal region was primarily disordered and had low predicted scores (pLDDT<60) (Fig. 4B). Transcription factors typically are disordered (Liu *et al*. 2006), and this was especially noted for plant MYB factors (Millard *et al*. 2019). DISOPRED3 and IUPred3 analyses of Z MYB and its orthlogs found the proteins to be primarily disordered except for the MYB regions (Fig. 4B). The AF2 prediction found the four amino acids essential for interaction with a target promoter in the Arabidopsis WER MYB protein (Pv Z residues: K51, N102, K105, N106) (Wang, 2020) are appropriately positioned on the recognition helices of the R2 and R3 MYB domains. Further, the DxxxLxxRLxxLx_13_R motif, shown to be responsible for the interaction between At WER and its bHLH partner, EGL3, is located in helix 1 of the R3 MYB domain (Wang, 2020). This, along with the sequence conservation and phylogenetic analysis, supports the role of Z MYB in the regulation of flavonoid production.

The P bHLH (Pv5-593.07G170300) protein contains a MYB-interacting region (MIR), a helix-loop-helix (HLH), and ACT-like domains typical of bHLH proteins associated with regulation of flavonoid LBGs (McClean *et al*. 2018). Its AF2 model (Supplementary Fig. S5) identified three well folded domains: the N-terminal MIR domain (residues 5-190) involved in bHLH/MYB protein interactions, a bHLH domain consisting of the basic residues 456-471, the HLH (residues 472-528) required for homo- or hetero dimerization with bHLH proteins (Feller *et al*. 2011), and the C-terminal ACT-like domain (residues 578-656), which is also involved in dimerization (Feller *et al*. 2006). The ACT-like domain has the characteristic ββαββα fold structure associated with plant bHLH proteins that regulate flavonoid biosynthetic genes (Kong *et al*. 2012). The conserved HER domain associated with binding of the protein to target promoters (Feller *et al*. 2011) is located in the basic region (463-471). All three domains are predicted with high confidence (pLDDT>70) (Fig. 4C). The rest of the protein was disordered with low confidence (pLDDT < 50). The DISOPRED3 and IUPred3 analyses agree with the AF2 prediction of disorder for residues between the folded domains (Fig. 4C). It was previously shown that protein MYB functions as a master regulator of flavonoid expression in seed coats and was predicted to be part of the common bean MBW complex (McClean, *et al*., 2018).

The AF2 T WDR, Z MYB, and P bHLH AF2 structure predictions were compared with similar proteins found in the protein databank (rcsb.org). The RMSD value between T WDR domain and the WDR domain of COP1 (At2g32950; PDB id: 6QTV), a seven blade, four WD-40 domain protein that acts as a photomorphogenesis repressor (Lau and Deng 2012), was 1.219 Å (over 159 residues). The RMSD value between the Z MYB R2R3-MYB domains and the crystal structure of the R2R3-MYB domains of the WER (At5g14750; PDB id: 6KKS) that regulates epidermal cell fate (Lee and Schiefelbein 1999) was 0.827 Å (over 99 residues). Lastly, RMSD value between the P MIR domain prediction and the N-terminal domain of Enhancer of Glabra 3 (EGL3) (At1g63659; PDB id: 7FDN) was 0.780 Å over 148 residues. EGL3 is the bHLH protein component of the Arabidopsis MBW complex regulating flavonoid biosynthesis (Gonzalez *et al*. 2008). Thus, the AF2 models are in good agreement with structures of known homologs.

### Structural prediction for mutant T and Z alleles gene products

t^65-73^ WDR lacks N-terminal residues 9-21 of the T WDR. Yet, AF2 predicts this protein still forms a seven β-propeller structure like T WDR (Fig. 4A). The key difference in the prediction for this mutant was the compensatory replacement of the d β-sheet of the seventh propeller with T WDR residues S3, T4, and Q5. Superposition of the t^65-73^ WDR model on the T WDR model gave an RMSD value of 0. 674 Å (over 300 residues). Thus, it is likely there is little impact on protein stability as well on any potential binding surfaces needed to function as the scaffold of the MBW complex.

The single base pair deletion of *t*^EW^ introduces an early stop codon after residue 254. The AF2 structure of t^EW^ WDR lacks the sixth and seventh blades from the β-propeller. Despite being able to form five blades, the t^EW^ WDR prediction did not close the propeller. Superposition of the *t*^EW^ model on the T WDR model gave an RMSD value 0. 674 Å (over 254 residues). The loss of these two blades could have a significant impact on protein stability as well as potential binding surfaces needed to bind the βHLH or MYB components of the MBW complex.

A K105N amino acid substitution distinguished the mutant z^EW^ MYB AF2 structure from Z MYB (Supplementary Fig. S4). The lysine residue, invariant at this position across the phylogeny of functional MYB proteins associated with flavonoid biosynthesis (Fig. 2B), is critical in DNA binding and mutating this residue in the WER protein “dramatically decreased” the interaction with its target promoter (Wang*et al*. 2020). The second difference is a four-residue deletion, Q178-E181 found in the unstructured region of z^EW^ MYB.

### Computational modeling of a common bean MBW complex

Previous research (McClean *et al*. 2018), and the results presented here, strongly suggest that the *Z*, *P*, and *T* genes encode the MYB, bHLH, WDR proteins of a plant MBW complex associated with proanthocyanin biosynthesis. Therefore, AF-Multimer was used to predict how these components could possibly assemble. To increase the potential reliability of these predictions, only residues with pLDDT values >70 for the three proteins were considered. We compared each protein structure modeled within the complex prediction to the corresponding monomer prediction. The structural predictions of Z MYB, P bHLH, and T WDR in the complex agreed well with the predictions of the monomer [RMSD = 1.077 (over 74 residues), 0.508 (over 104 residues), and 0.397 (over 336 residues), respectively]. We also predicted the soybean and Arabidopsis MBW protein complexes with AF-Multimer and compared each protein component in the modeled complex predictions to the corresponding individual protein predictions. For the soybean MBW, the GmTT2 MYB, GmTT8 bHLH, and GmTTG1 WDR proteins, the RMSD values were 0.996 (over 59 residues), 1.192 (over 84 residues), and 0.368 (over 301 residues, respectively. For the Arabidopsis MBW complex, RMSD values for the AtTT2 MYB, AtTT8 bHLH, and AtTTG1 WDR proteins RMSD = 1.016 (over 7 residues), 0.474 (over 179 residues), and 0.483 (over 323 residues), respectively. (n=224)

The modeled Z MYB/P bHLH/T WDR MBW complex was analyzed with the PISA server to identify interacting regions. The Z MYB and T WDR interaction involves two β-strands formed by Z MYB residues 147-149 and 151-154, as well as strands βP3d (residues 161-167) and βP4d (residues 204-209) of T WDR (Fig. 5A). The two Z MYB β-strands potentially define a β-MoRF, an intrinsically disordered region in the unbound T MYB that transitions to a β structure in the presence of a binding partner (Mohan *et al*. 2006). A similar structural transition to a β structure was also found for the homologous residues in the predicted soybean GmTT2 MYB and Arabidopsis AtTT2 MYB proteins only in their corresponding MBW complexes. The motif defined by the two β-strands of Z MYB is only found among MYB proteins experimentally confirmed to regulate PA synthesis (Fig. 3B). This provides additional support of a potential function role of the motif in the binding MYB to WDR proteins in MBW complexes associated with PA biosynthesis. This Z MYB and T WDR interface accounts for 2.2% of the solvent accessible area (SAA) of Z MYB and 4.0% of the SAA of T WDR. The interaction is stabilized via hydrogen bonds involving Z MYB residues V148, T150, K151, and T153 and residues Q165, I167, E202, S204, and I206 of T WDR.

**Fig. 3.**
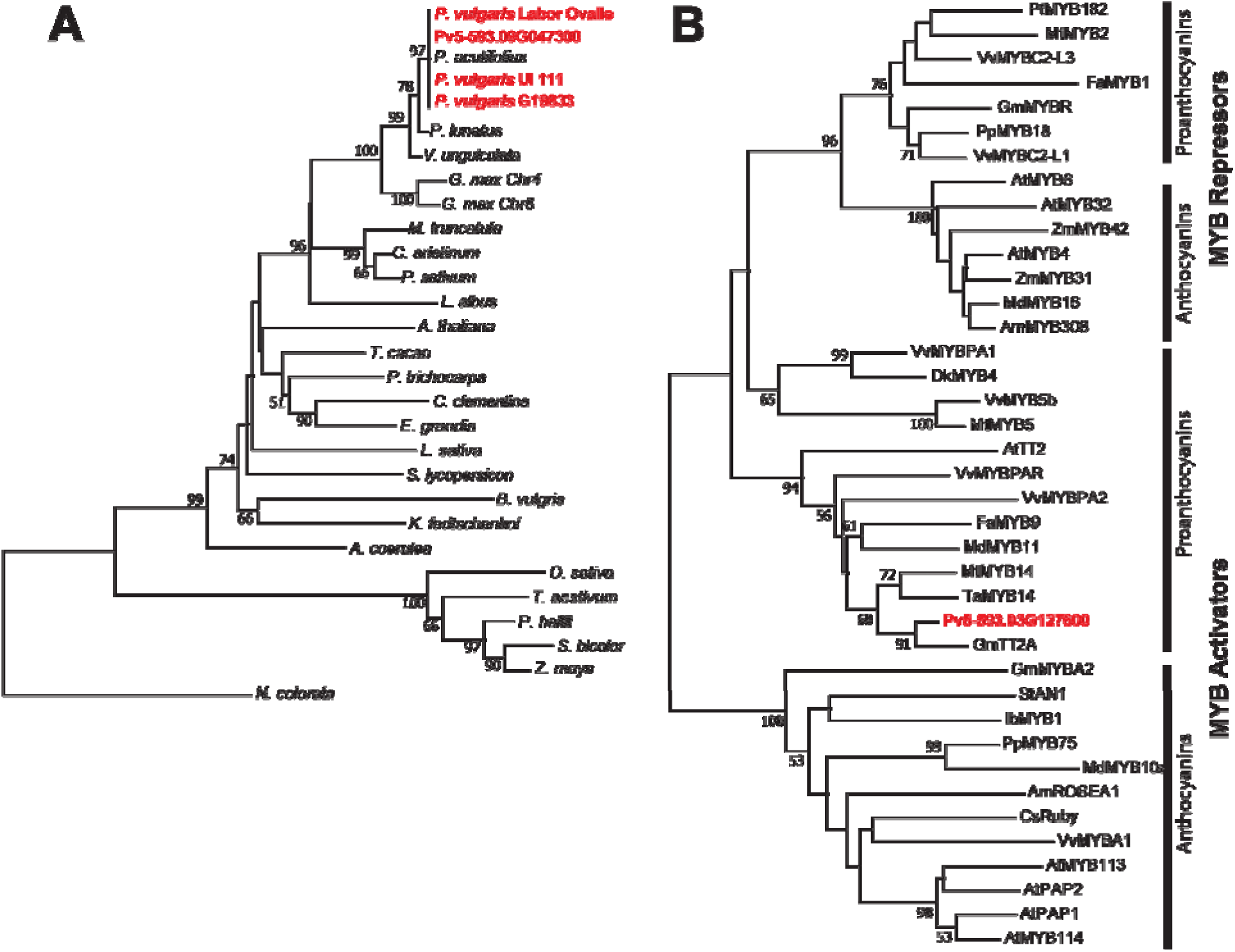
Neighboring joining trees of WDR homologs of common bean (A) angiosperm WDR proteins and (B) MYB proteins experimentally to act as repressors or activators of flavonoid biosynthesis. Common bean gene models are shown in red.

**Fig. 4.**
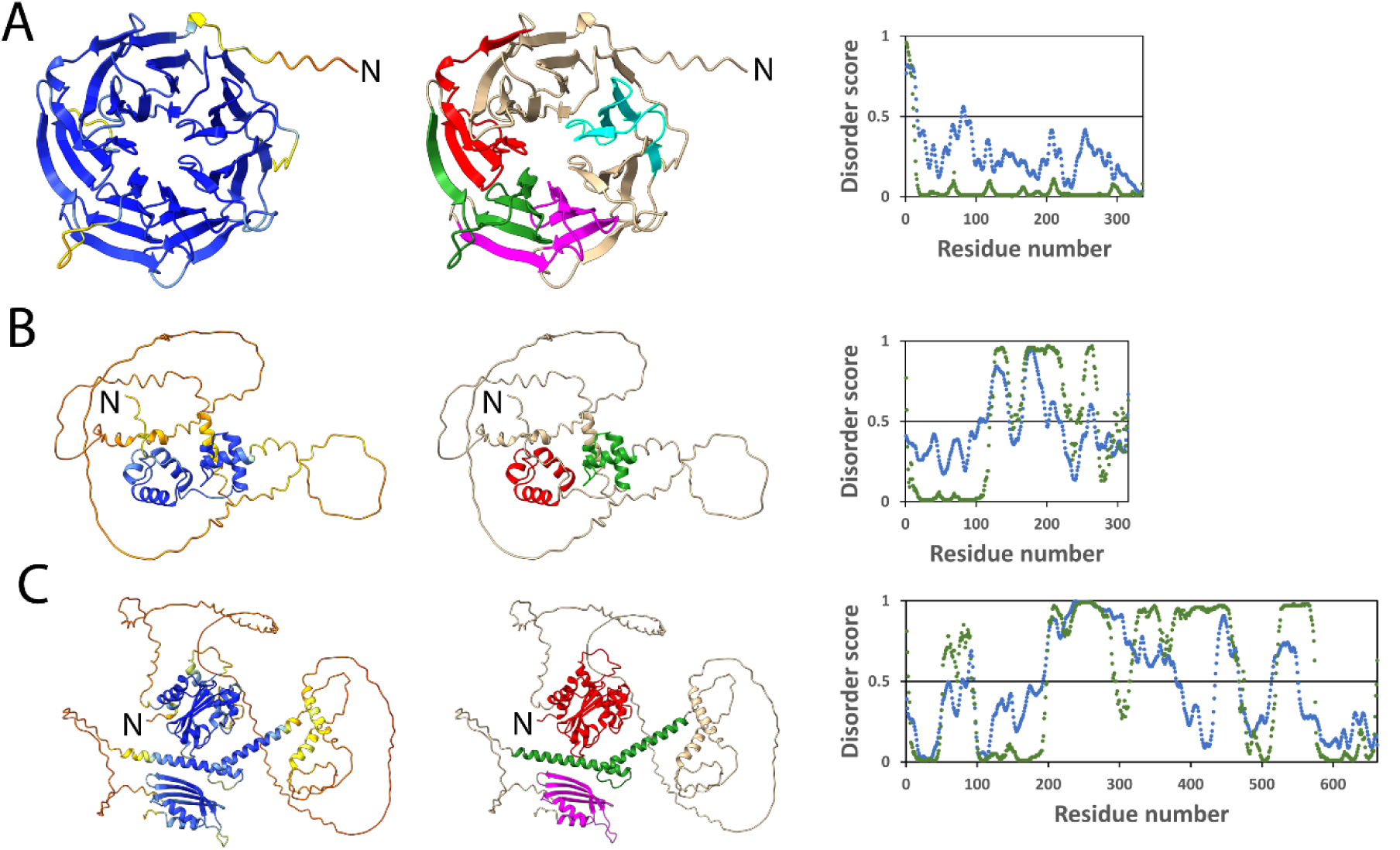
Structural details of the AlphaFold2 protein models of the (A) *T* WDR, (B) *Z* MYB, and (C) *P* bHLH genes. For each protein the left model is the pLDDT values across the protein. High pLDDT values for a residue suggests a good structural fit. Blue represents high confidence, yellow is moderate confidence, and red and orange is poor confidence. The central figures show the location of the domains in the proteins. For *T* WDR protein (A), the four WD40 domains are depicted. The R2 and R3 MYB domains are shown for the *Z* MYB protein (B). The MYB-interacting region, helix-loop-helix, and activation domains are highlighted for *P* bHLH protein. Note that these domains were modeled with high confidence based on their pLDDT values. The right most image for each protein is the disorder score. The green represents the DISOPRED3 score, and the blue represents the IUPred3 score. Regions with disorder scores >0.5 are considered to be disordered. Note that the disorder regions also have low pLDDT values.

**Fig. 5.**
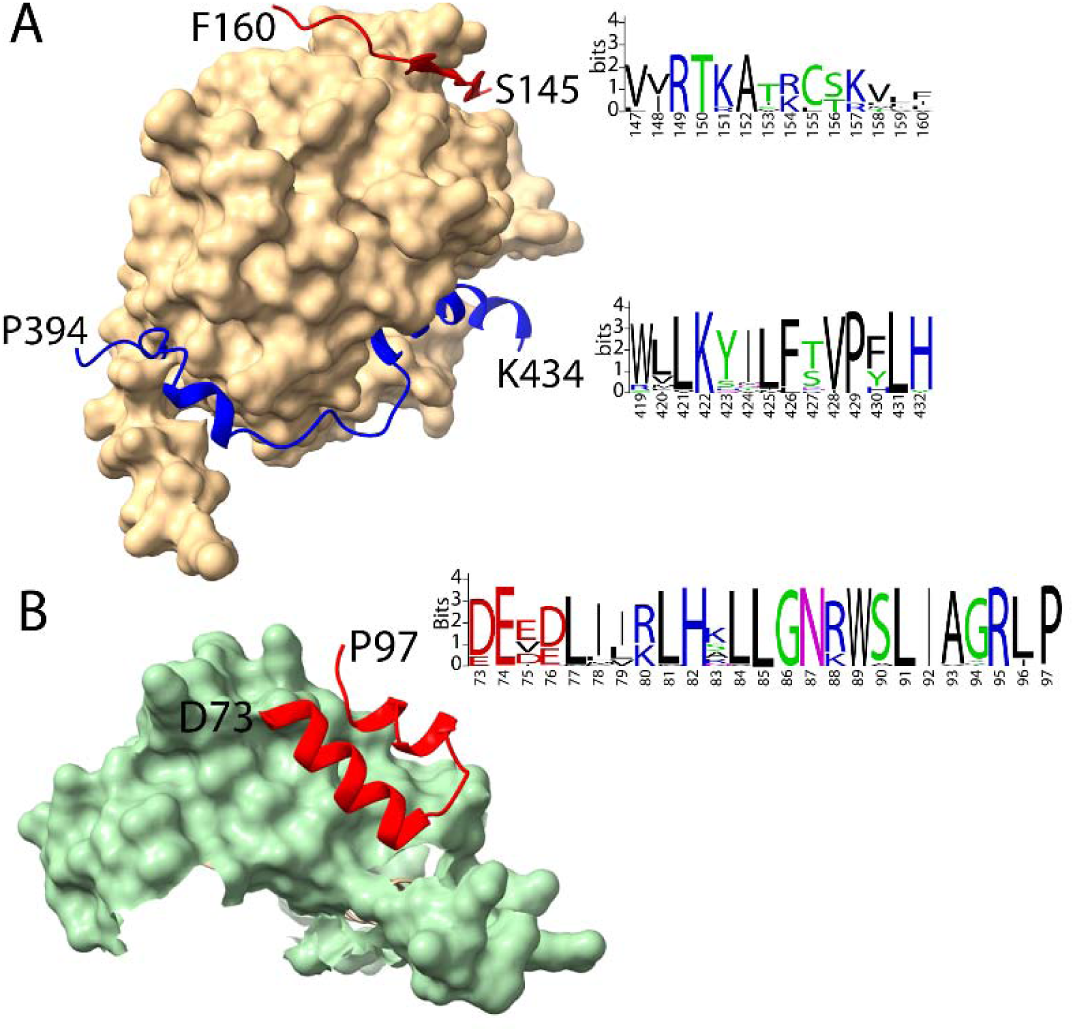
Interfaces between components of an AlphaFold-Multimer predicted model of a MBW complex of common bean. (A) The interfaces between the T WDR protein (tan) and its interfacing regions with the Z MYB (red) and P bHLH (blue) proteins. To the right are sequences of high conservation found in other orthologs of the Z and P proteins. (B) The interface between the MIR region of the P bHLH protein (green) and the Z MYB R3 domain (red). To the right is the sequence of a highly conserved domain found among homologs of the Z protein.

In the predicted MBW complex, P bHLH residues 395-431 wrap around T WDR starting at the propeller βP5-βP6 interface and across the top of toroid axis and disengaging at the top of the βP2-βP3 interface (Fig. 5A). PISA analysis of this interaction indicates the P bHLH interface accounts for 11.7% of the SAA of the protein, while the SAA of the T WDR protein interface accounts for 10.1% of its total SAA. The contact residues of P bHLH with T WDR include two helices formed by residues 404-408 and residues 419-431. These two regions were also observed between GmTT8 bHLH and GmWD40 WDR in the predicted soybean MBW complex. The 419-432 region of P MYB contains highly conserved sequences among bHLHs that are components of other plant MBW complexes (Fig. 5A). This region is well a supported helical structure with pLDDT values >70.

The interface between Z MYB and P bHLH involves Z MYB residues R3α1 and R3α2 within the R3 domain which leaves R3α3 residues free to bind to DNA (Fig. 5B, Fig. S2). P bHLH provides residues helices α2, α3, and α5 within its predicted MIR domain for the interaction (Fig. 5B, Fig. S5). Validation of the predicted model comes from its alignment with the interface found in the co-crystal structure of the At WER MYB and At EGL3 bHLH proteins (PDBID: 7FDL; Wang et 2022). The Z MYB and P bHLH interaction accounts for 4.0% of the Z MYB SAA and 2.1% of the P bHLH SAA. The interface regions contain two salt bridges: R88 of Z MYB and E98 of P bHLH, and R95 of Z MYB and E 102 of P bHLH. The Z interface involves the highly conserved R3 domain, which has a similar structure in all experimentally determined flavonoid-activating MYB proteins that interact with a homolog of P bHLH. Ten Z MYB R3 residues are invariant in the MYB homologs. Of the Z MYB residues predicted to interact with P bHLH, five (E74, L85, L91, R95, P97) are invariant in Z MYB orthologs (Fig. 5B).

### Modeling of the P bHLH homodimer and Z MYB/P bHLH heterodimer

Since transcription factors, including bHLHs, are known to function as dimers (Murre *et al*. 1989), AF-Multimer was used to predict the homodimeric structure of the P protein. AF-Multimer enforces a global C2 symmetry axis. Therefore, all dimeric interfaces were placed along a single C2 axis. However, the large, disordered regions between each of the dimer interfaces do not structurally predicate that there is a single global axis, but more likely, three individual local 2-fold symmetry axes. Thus, the individual interfaces are likely valid, but their location in three-dimensional space relative to each other is not. For example, the classic bHLH dimer is defined by a helical region that binds DNA followed by a leucine zipper that forms the dimer interface. P bHLH residues 459-484 bind DNA and residues 493-528 contain the leucine zipper (Fig. S5). However, a second dimer interaction between symmetrically related helical residues 416-432 is modeled in the DNA binding space, which would preclude the function of DNA binding. Interactions between ACT-like domains have been shown to mediate bHLH homodimer formation (Feller *et al*. 2006). Indeed, a third dimer interface was predicted to be formed between the two ACT-like domains. This interface is similar to that found for other ACT domains such as in *E. coli* phosphoglycerate dehydrogenases [PDB IDs: 1PSD, (Schuller *et al*. 1995); 1YBA (Thompson *et al*. 2005)]. Further support of this interface is that P bHLH residue V615, which is buried at this interface, corresponds positionally to a critical residue that determines the strength of the ACT-like domain interaction (Lee *et al*. 2021). However, the position of the ACT-like domains relative to the rest of the complex is defined by this interface residing on the global C2 axis. Therefore, while each interface may exist, the local relationships of the individual C2 axes is likely asymmetric and reliable conclusions are difficult to discern relative to the orientations of the domains.

Since plant MYB and bHLH proteins involved in flavonoid expression are known to interact as dimers (Goff *et al*. 1992), the Z MYB and P bHLH proteins were also modeled as a dimer. The interfaces regions of the two proteins modeled as a dimer were the same as seen for the full MBW complex.

### Effects of mutant Z MYB and T WDR proteins on common bean MBW complex interactions

Protein interactions with MBW complexes containing mutant z^EW^ MYB or t^EW^ WDR were evaluated using PISA. The model predicted MYB R3 helix 3 residues 107-115 of z^EW^ MYB interact with WD40 domain 2 residues 119-127 T WDR as part of the mutant but not the wild type MBW complex. While both of these regions are well predicted (80> pLDDT >60), closer examination of this interface reveals the presence of several severe steric clashes that preclude this interface from being real. Therefore, it is unsurprising that this interaction is not observed in the wild type Z MYB/P bHLH/T WDR complex. However, the z^EW^ K105N substitution would severely disrupt z^EW^ from interacting with its target promoter. In the crystal structure of WT At WER MYB complexed with DNA (PDB ID:6KKS), K105 forms extensive interactions in the major groove of the DNA (Wang et al. 2020). The equivalent residue in z^EW^, N105, would lack these major groove interactions. Wang et al. (2020) found this MYB region was important to promoter binding. The interaction between P bHLH and T WDR in the presence of z^EW^ remains largely unchanged from that predicted for the wild type complex where P bHLH residues 395 – 431 wrapped around T WDR. However, there is an additional helical region, defined by P bHLH residues 361-379, that is modeled with high confidence (92> pLDDT >72). This region interacts with propeller strands βP4 βP5 of T WDR and completes an offset belt that wraps around the T toroid.

A major structural effect of the C-terminal deletion of t^EW^ WDR was the elimination of the sixth and seventh blades from the β-propeller, thus forming an unclosed five bladed structure. While five-bladed propellers are possible, the inability of t^EW^ WDR to close leaves exposed hydrophobic surfaces that provide a strong driving force for false protein interfaces. Indeed, the ACT-like domain from P bHLH tries to extend the β-structure at the edge of propeller βP5. While this interface is modeled with high confidence, it likely relies on the extended beta structure. Before any reliable conclusions can be drawn from this predicted model, experiments would need to be performed to validate these non-physiological interfaces.

## Conclusion

While multiple common bean genes (*G*, *B*, *V*, *R*, *Rk*, *Sal*) control seed coat color, *T* and *Z* control the spatial pattern of seed coat color expression. This led to the working hypothesis that both genes were regulators of the flavonoid biosynthetic pathway. Physical intervals containing*T* and *Z* were determined here based on previous marker data, and multiple analyses supported the conclusion the *T* encodes a WDR protein and *T* encodes a MYB protein and that these genes are critical to the expression of flavonoids in common bean. Domain and protein modeling found the *Z* gene candidate encoded a mostly unstructured R2R3 MYB protein, while the *T* gene candidate encoded a highly structured, seven blade WDR protein with four WD40 domains. Phylogenetic analysis found these two proteins clustered with other WDR and MYB proteins experimentally shown to regulate genes of the flavonoid pathway. AF2 models of these proteins. In addition, the previously cloned P bHLH protein was found to be highly similar to other bHLH proteins that function as a component of other defined plant MBW complexes (Payne *et al*. 2000; Zhang *et al*. 2003).

Experimental structural models of MBW complex proteins are rare. Only a single model consisting of the WER R2R3 MYB (Wang *et al*. 2020) and EGL3 bHLH proteins (Wang et al. 2022) has been reported. No experimental model of a WDR protein that functions in a MBW complex is known. RMSD analyses of AF2 models of Z MYB and P bHLH found proteins had good structural fits with WER and EGL3, respectively, that function in an Arabidopsis MBW complex. Since no full MBW complex has been described, a model with bean Z MYB, P bHLH, and T WDR proteins was predicted with AF2-multimer. The complex interface of Z MYB and T WDR involved a potential β-MoRF in a Z MYB motif that is unstructured in the monomer state. This motif is conserved among plant proteins that regulate PA biosynthesis. The P bHLH and T WDR interface involves a structured region that is conserved among other bHLH proteins that function in a MBW complex. The Z MYB and P bHLH interact with different regions of T WDR in bean, soybean, and Arabidopsis complexes. Finally, Z MYB and P bHLH interact in the same regions of each protein as predicted from crystal structure analysis of Arabidopsis WER and EGL3 (Wang *et al*. 2022). Structures were also predicted for the bean MBW complex containing mutant t^EW^ WDR or z^EW^ MYB proteins. In the t^EW^ WDR complex, its interaction with P bHLH was compromised by the deletion of the βP6 blade, which resulted in P binding elsewhere to T WDR. The introduction of the z^EW^ MYB protein in the complex created a new interaction that would comprise the ability of z^EW^ MYB to function. These predictions can act as hypotheses for further crystal structural analyses.

## Supplementary Data

The following supplementary data are available at *JXB* online.

**Supplementary Fig.S1.**
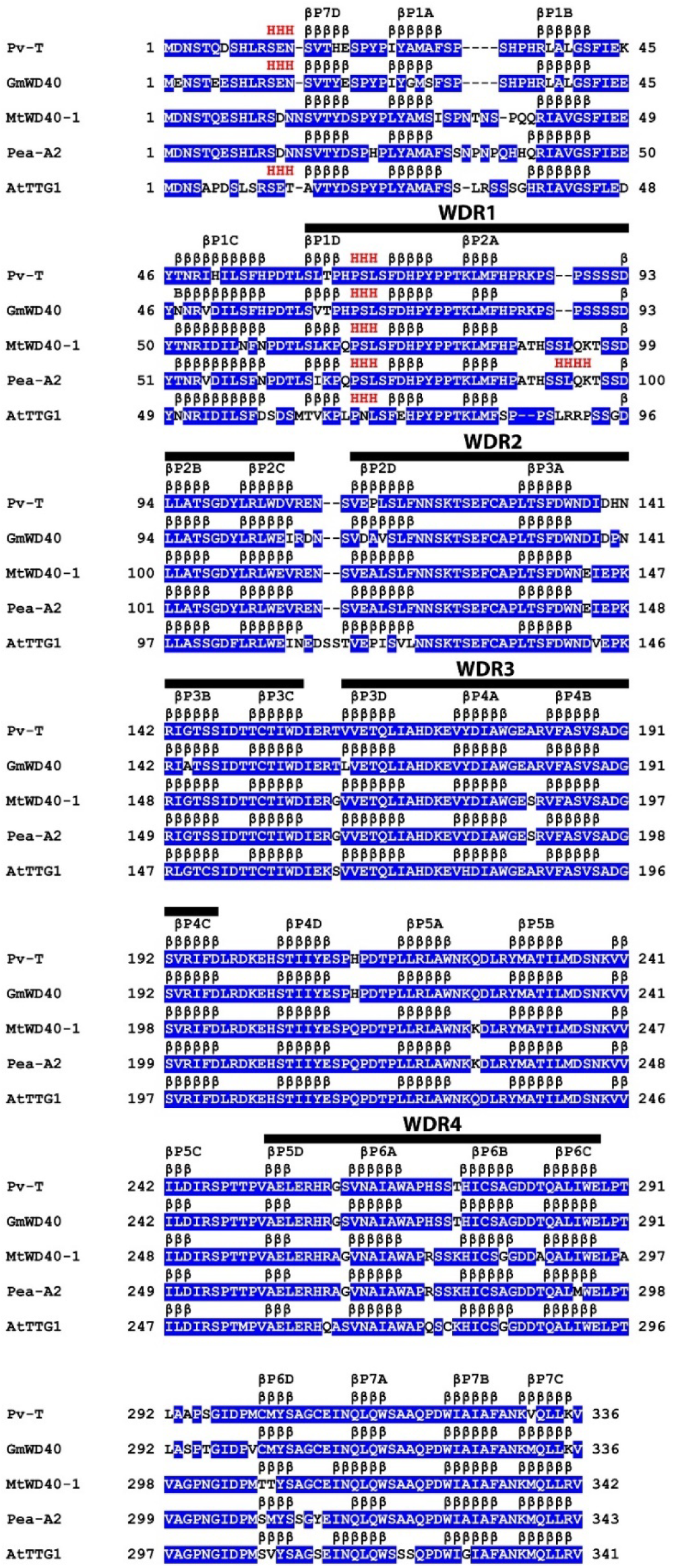
Multiple sequence alignment of WDR proteins of common bean *T* (Pv5-593.09G047300), soybean GmWD40, *M. truncatula* MtWD40-1, pea A2, and *A. thaliana* TTG1. The locations of α-helices and β-sheets, as predicted by AF2, are designated as consecutive “H”s and “β”s. Each β-propeller blade is designated βP#, and each β-sheet within a blade is designated a, b, c, or d. The position of each WD-40 repeat (WDR) is shown.

**Supplementary Fig. S2.**
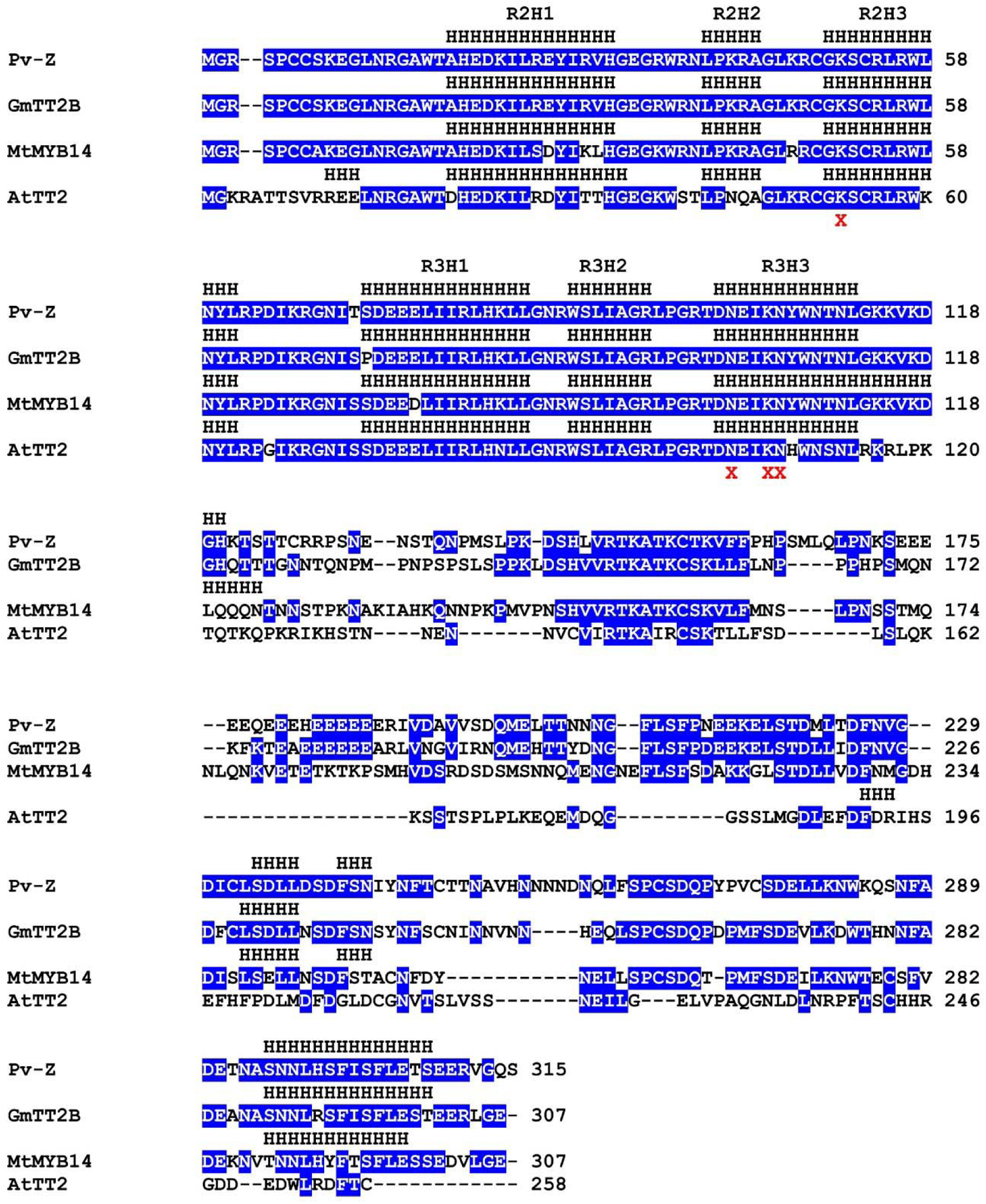
Multiple sequence alignment of common bean *Z* (Pv5-593.03G127600), soybean GmTT2B, *M. truncatula* MtMYB14, and *A. thaliana* AtTT2. The locations of α-helices, as predicted by AF2, are designated as consecutive “H”s. R2 and R3 designated the two MYB domains, and H1, H2, and H3 designated the helices in each of the MYB domains. Conserved Arabidopsis MYB domain residues experimentally shown to interact with DNA of a target promoter (Wang *et al*. 2020) are shown with an X.

**Supplementary Fig. S3.**
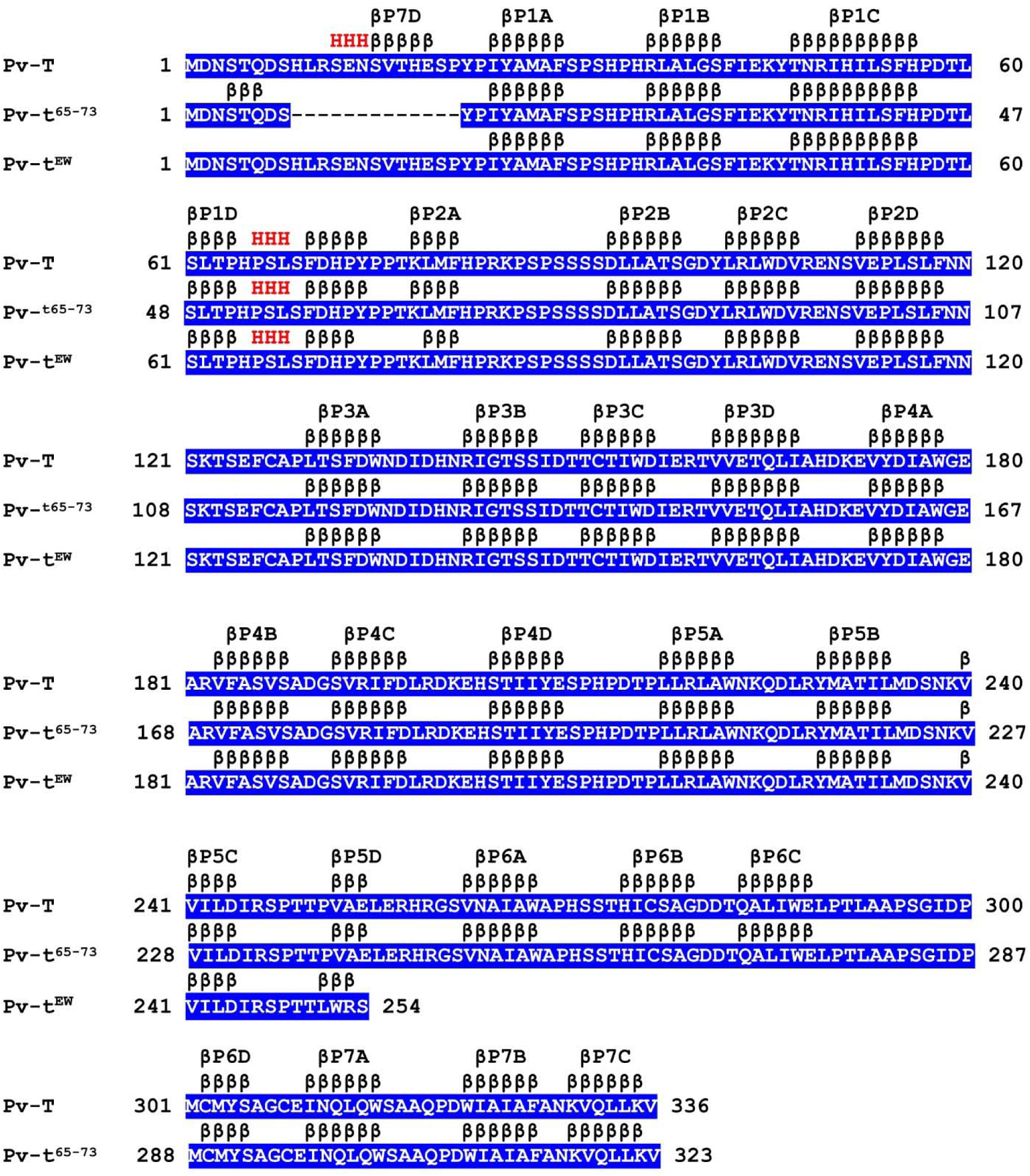
Structure of wild type (Pv-T) and mutant (Pv-t^65-73^ and Pv-t^EW^) proteins. The mutant proteins are associated with partial seed color and white flowers. The locations of α-helices and β-sheets, as predicted by AF2, are designated as consecutive “H”s and “β”s. Each β-propeller blade is designated βP#, and each β-sheet with a blade is designated a, b, c, or d.

**Supplementary Fig. S4.**
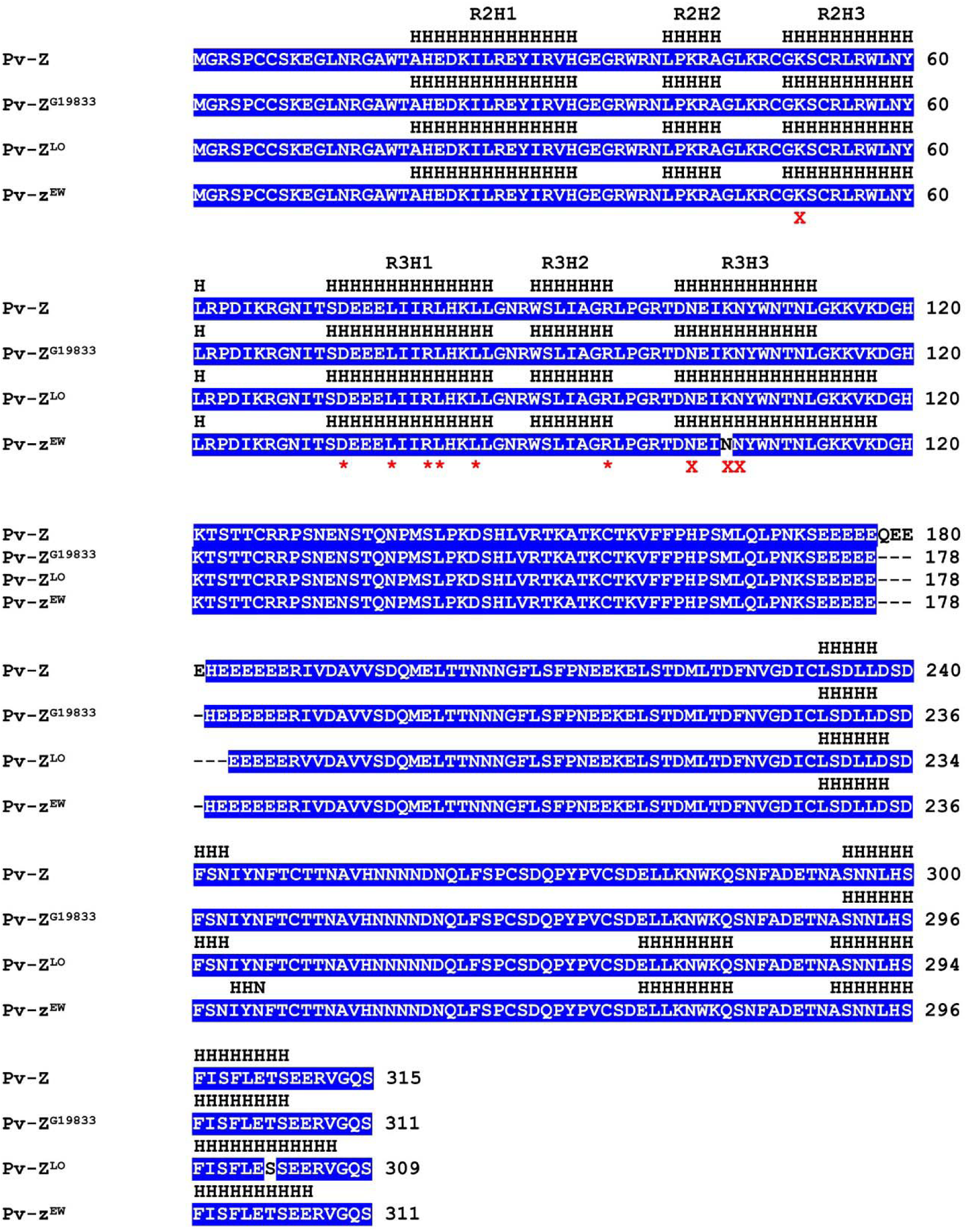
Structure of common bean wild type (Pv-Z, Pv-Z^G19833^, and Pv-Z^LO^) and mutant (Pv-z^EW^) proteins. The locations of α-helices, as predicted by AF2, are designated as consecutive “H”s. R2 and R3 designate the two MYB domains, and H1, H2, and H3 designate the helices in each of the MYB domains. Conserved MYB domain residues experimentally shown to interact with DNA of a target promoter (Wang *et al*. 2020) are shown with an X. Conserved residues that interact with a bHLH partner are shown with an * (Wang *et al*. 2022).

**Supplementary Fig. S5.**
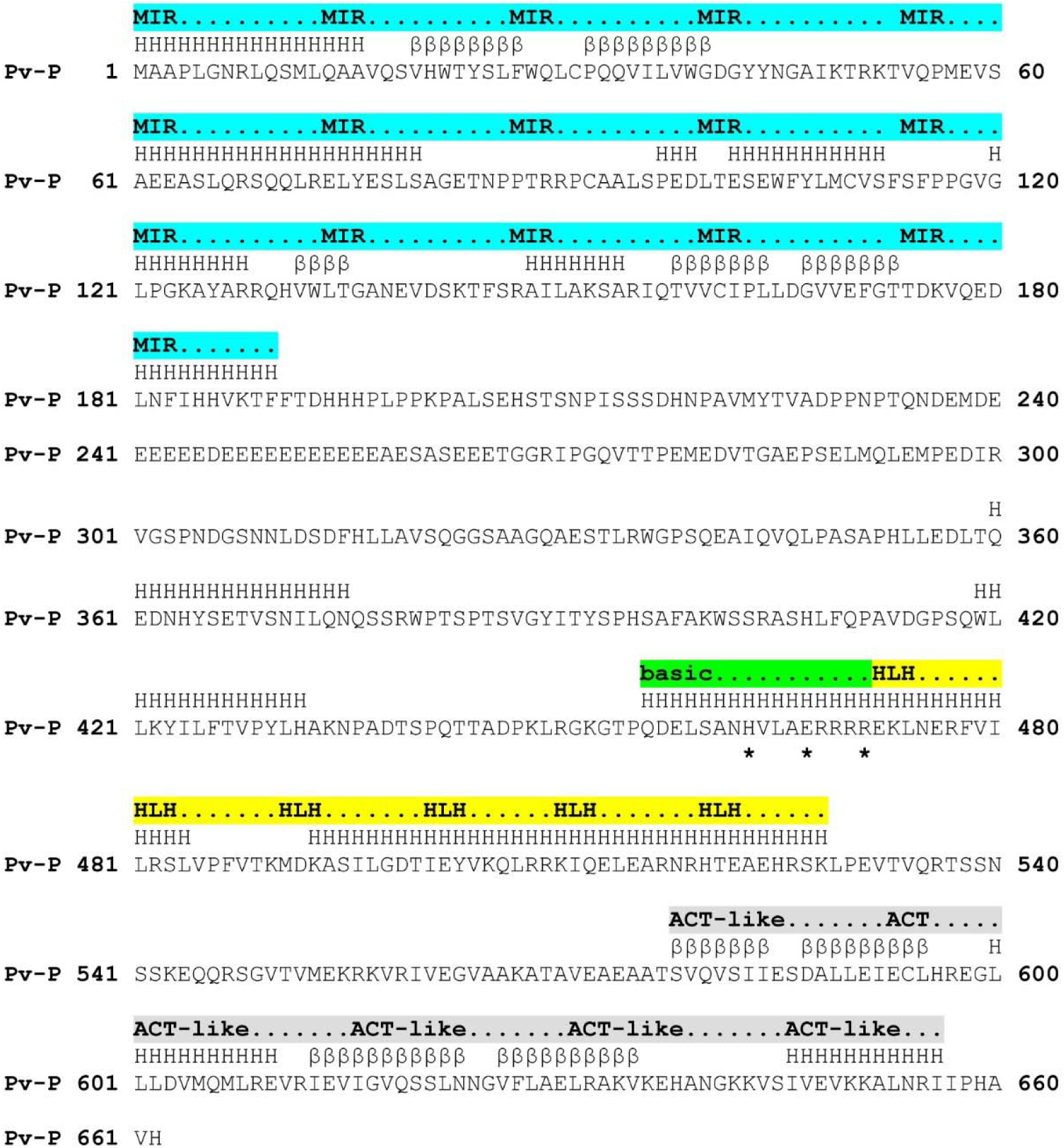
Common bean wild type P protein sequence, domain boundaries and AlphaFold2 predicted secondary structure elements. The MYB-MIR-interacting region (MIR) is highlighted in cyan, the bHLH basic region highlighted in green, the helix-loop-helix (HLH) domain is highlighted in yellow, and the activation domain (ACT) is highlighted in grey. The three asterisks below the basic region are the Hx_3_Ex_3_R motif associated with binding to target promoters. Secondary structure elements are indicated above the sequence with residues predicted to form helical structure designated by Hs, and residues predicted to form beta strands notated by consecutive β symbols.

Table S1. Select Bassett introgression lines genotyped with the tEW and t65-73 PACE markers.

Table S2. T and Z genotypes of members of the Middle American Diversity Panel.

Table S3. T and Z genotypes of members of the Andean Diversity Panel.

Table S4. Sequences of primers used to amplify amplicons from the T gene.

Table S5. The sequences of the PACE primers used to genotype the tEW and t65-73 alleles of the T gene.

Table S6. T and Z populations used to map candidate genes.

Table S7. Sequence IDs of a representative sample of higher plant T protein orthologs used for the phylogenetic analysis. The E-value and % identity values are from a blastp analysis using Pv5-593.09G047300 query against each of the Phytozome reference genomes to identify the corresponding ortholog,

Table S8. Protein sequences used for the phylogenetic analysis of Z gene candidate (Pv5-593.03G127600) and proteins known to act as activators or repressors of proanthocyanin or anthocyanin biosynthesis. The E-value and % identity values are from a blastp analysis using Pv5-593.03G127600 as a query against the MYB proteins to calculate E-value and percent identity.

## Authors contributions

PEM, PNM, JMP: conceptualization; RL: data curation; PEM, JR, CLC, CO, RL: formal analysis; PEM, PNM, JMO: funding acquisition; JR, CLC, CO, RL: Methodology; PEM, CCLC: visualization; PEM, JR, CLC, CO, RL, PNM, JMO: writing – review and editing.

## Conflict of Interest

The authors declare no conflicts of interest.

## Funding

Funding for this research was provided by the USDA, Agricultural Research Service through the Pulse Crop Health Initiative, Agreement no. 58-3060-0-041.

## Data availability

The data supporting this research can be found within the main body of the paper or the supplementary data published online.

## Notes

### Competing Interest Statement

The authors have declared no competing interest.

